# Deciphering D4Z4 CpG methylation gradients in fascioscapulohumeral muscular dystrophy using nanopore sequencing

**DOI:** 10.1101/2023.02.17.528868

**Authors:** Russell J Butterfield, Diane M Dunn, Brett Duval, Sarah Moldt, Robert B Weiss

## Abstract

Fascioscapulohumeral muscular dystrophy (FSHD) is caused by a unique genetic mechanism that relies on contraction and hypomethylation of the D4Z4 macrosatellite array on the chromosome 4q telomere allowing ectopic expression of the *DUX4* gene in skeletal muscle. Genetic analysis is difficult due to the large size and repetitive nature of the array, a nearly identical array on the 10q telomere, and the presence of divergent D4Z4 arrays scattered throughout the genome. Here, we combine nanopore long-read sequencing with Cas9-targeted enrichment of 4q and 10q D4Z4 arrays for comprehensive genetic analysis including determination of the length of the 4q and 10q D4Z4 arrays with base-pair resolution. In the same assay, we differentiate 4q from 10q telomeric sequences, determine A/B haplotype, identify paralogous D4Z4 sequences elsewhere in the genome, and estimate methylation for all CpGs in the array. Asymmetric, length-dependent methylation gradients were observed in the 4q and 10q D4Z4 arrays that reach a hypermethylation point at approximately 10 D4Z4 repeat units, consistent with the known threshold of pathogenic D4Z4 contractions. High resolution analysis of individual D4Z4 repeat methylation revealed areas of low methylation near the CTCF/insulator region and areas of high methylation immediately preceding the *DUX4* transcriptional start site. Within the *DUX4* exons, we observed a waxing/waning methylation pattern with a 180-nucleotide periodicity, consistent with phased nucleosomes. Targeted nanopore sequencing complements recently developed molecular combing and optical mapping approaches to genetic analysis for FSHD by adding precision of the length measurement, base-pair resolution sequencing, and quantitative methylation analysis.

## INTRODUCTION

Fascioscapulohumeral muscular dystrophy (FSHD) is one of the most common forms of muscular dystrophy with a prevalence of 1:15,000-1:20,000 (Padberg 1982; Flanigan et al. 2001; Mostacciuolo et al. 2009). As suggested in the name, FSHD results in weakness of the muscles of the face, shoulder stabilizers, and upper arm but also affects proximal and distal muscles of the legs. Unlike other muscular dystrophies, weakness can be asymmetric, and onset and severity are variable even between members of the same family with the same mutation (Tyler and Stephens 1950; Flanigan et al. 2001). Weakness is progressive, leading to significant morbidity with ∼20% requiring a wheelchair over 50 years of age, and 10% developing restrictive respiratory disease (Scully et al. 2014; Statland and Tawil 2014). Complicating diagnosis, FSHD has a mechanism combining both genetic and epigenetic factors in a region that is notoriously difficult to analyze due to its length and repetitive elements.

FSHD maps to the subtelomeric end of chromosome 4q, in a region that contains tandem copies of D4Z4, a 3.3kb CpG-rich macrosatellite with an intronless retrogene encoding the DUX4 homeobox domain protein. *DUX4* is an important regulator of early genome activation, but expression is heavily repressed by epigenetic mechanisms after the first few cell divisions (Hendrickson et al. 2017). In FSHD patients, chromatin relaxation accompanied by hypomethylation of the D4Z4 array leads to ectopic expression of *DUX4* which is toxic to skeletal muscle (Lemmers et al. 2010a). Hypomethylation of the array is asymmetric, involving the proximal more than distal D4Z4 repeats, and is inversely correlated with the number of repeats (de Greef et al. 2009). *DUX4* expression in skeletal muscle arises from the distal D4Z4 repeat and requires a short flanking sequence, pLAM, which contains a 3’-terminal exon and polymorphic signals for pre-mRNA cleavage and polyadenylation (Lemmers et al. 2007; de Greef et al. 2009). pLAM and related polymorphic sequences are found in FSHD permissive 4qA haplotypes. Non-FSHD permissive haplotypes include the nearly identical sequences on chromosome 10q which lack the polyadenylation signal, 4qA haplotypes that have pLAM but lack the polyadenylation signal, and 4qB haplotypes which lack pLAM altogether. These non-permissive haplotypes are not known to result in FSHD despite hypomethylation of the contracted array (Lemmers et al. 2010a).

Most individuals have 11-100 *D4Z4* repeats in the array, but contraction of the array to 10 or less copies on a permissive 4qA haplotype results in FSHD1 (MIM158900) (Lemmers et al. 2010a). FSHD1 is the most common form of FSHD, accounting for 95% of cases (Rieken et al. 2021). Contraction length of the *D4Z4* array is correlated with disease severity but does not explain all the variability. Individuals with 1-3 *D4Z4* repeats usually have severe disease, individuals with 4-7 repeats typically have intermediate disease manifestations, and individuals with 8-10 repeats are usually mildly affected (Salort-Campana et al. 2012; Statland and Tawil 2014; Statland et al. 2015). FSHD2 (MIM 158901) is phenotypically identical to FSHD1, but has a digenic mechanism involving both a permissive 4qA haplotype and mutation in one of several chromatin modifier genes (*SMCHD1*, *DNMT3B*, and *LRIF1*) (Lemmers et al. 2012; van den Boogaard et al. 2016; Sacconi et al. 2019; Hamanaka et al. 2020). The combined permissive 4qA haplotype and additional mutation in a chromatin modifier gene, results in hypomethylation of the D4Z4 array and expression of the *DUX4* gene, even without a full contraction of the array. However, most FSHD2 patients have 9-20 *D4Z4* repeats in the array suggesting that repeat length is an important factor in pathogenesis in addition to hypomethylation due to a mutation in the second gene (de Greef et al. 2009; Rieken et al. 2021).

Analysis of the FSHD-related D4Z4 array on 4q is complicated by the large size of the array, a nearly identical array on telomeric chromosome 10q, and the presence of divergent D4Z4 arrays scattered throughout the genome (Winokur et al. 1994; Lyle et al. 1995; Leidenroth et al. 2012). The gold-standard diagnostic test for FSHD1 is Southern blot after restriction digest which can differentiate the chromosome 4q and 10q *D4Z4* arrays and estimate their length (Zampatti et al. 2019). More recently, molecular combing and optical mapping techniques have emerged to estimate the size of the array (Vasale et al. 2015; Nguyen et al. 2017; Dai et al. 2020). These methods represent a significant advance, but like Southern blotting they are technically demanding and are only available at a few select laboratories. Neither method provides detailed information about the methylation state or background haplotype (4qA/4qB) without additional testing. Epigenetic analysis using targeted bisulfite sequencing can differentiate between FSHD and healthy controls using a direct measurement of methylation in sub-regions of the D4Z4 array and has been proposed as a novel diagnostic tool that does not require determination of the length of the array (Roche et al. 2019; Gould et al. 2021). Genetic testing is further complicated by relatively common translocations between the 4q and 10q arrays, duplication arrays, and somatic mosaicism (Rieken et al. 2021; Lemmers et al. 2022).

Emerging long-range sequencing technologies allow for sequencing across complex repetitive regions and address existing gaps in genetic analysis for FSHD. Recent publication of the first complete genome relied heavily on these new long-range sequencing tools to successfully sequence highly repetitive centromeric and telomeric sequences including both 4q, 10q, and other D4Z4 arrays (Nurk et al. 2022). Nanopore sequencing is a single-molecule long-range sequencing tool that can routinely produce read lengths of 30 kb or more (Deamer et al. 2016). Nanopore sequencing has been successfully used to sequence the 4q *D4Z4* array from a BAC clone, but an attempt to sequence the array from genomic DNA as part of a whole genome analysis resulted in only a small number of full-length reads from the *D4Z4* array limiting utility of this technique (Mitsuhashi et al. 2017). Nanopore Cas9-Targeted Sequencing (nCATS) is a recently developed tool to enrich target sequences for nanopore sequencing allowing marked increase in read depth for targeted sequences (Gilpatrick et al. 2020), and has been proposed as a novel tool to sequence the D4Z4 array in FSHD (Hiramuki et al. 2022). We used nCATS as a comprehensive tool for genetic analysis of FSHD to determine the length of the 4q and 10q D4Z4 arrays, differentiate the 4q from 10q arrays, determine A/B haplotype, identify divergent D4Z4 elsewhere in the genome, and estimate methylation across all CpGs in the array.

## RESULTS

### Targeted nanopore sequencing of D4Z4 repeat arrays

We used multiple guide RNAs (gRNAs) to target centromeric (p13E-11) and telomeric (pLAM) sequences flanking D4Z4 arrays on chromosomes 4qA and 10q (**Fig. 1A,B**). Human peripheral blood DNA was cut with a Cas9 p13E-11/pLAM gRNA mixture, followed by adapter ligation and nanopore sequencing on MinION flow cells. Targeted enrichment was observed at both 4q and 10q, as well as consistent enrichment peaks on other chromosomes with divergent D4Z4 arrays (**Fig. 1C**). We determined the size of the D4Z4 array in subjects with FSHD1 and FSHD2 with a range of 3-19 D4Z4 repeats on the 4q array (**Table 1, Supplemental Table S1**). The average 4q/10q on-target percentage of total reads was 0.7% from eight MinION libraries using the p13E-11/pLAM gRNA mixture (**Table 1, Supplemental Table S2**). FSHD1 participant P2 had a known 4qA 6U D4Z4 allele confirmed with multiple 17.5 kb, p13E-11 to pLAM, full length, cut-to-cut reads covering the full array (39 6U reads, **Table 1**, **Fig. 1D**). This enrichment was reproducible in participants P1, P3, and P4, with 50, 68, and 78 reads confirming the presence of their 4qA 6U D4Z4 alleles. Two additional FSHD1 participants had their pathogenic 4qA alleles confirmed, including an 8.0 kb 3U allele (P6) and compound heterozygous 9U alleles from participant P5, who was heterozygous for a 27.6 kb 4qAS 9U allele and a 29.0 kb 4qAL 9U allele. In FSHD2 participants P7 and P8, we identified 60.1 kb (19U) and 50.2 kb (16U) p13E-11 to pLAM reads spanning their shortest 4qA allele, respectively. In each of the eight samples, we identified reads that spanned 4qA or 10q D4Z4 arrays (**Table 1**). Full-length reads spanning shorter arrays were recovered at greater efficiency, consistent with a size bias operating at the nanopores. During the mapping of reads to the Telomere-to-Telomere (T2T) CHM13v2.0 reference genome, minimap2 was able to distinguish between highly similar 4q and 10q D4Z4 reads. This was demonstrated by the ratio of 4q- specific (*XapI*) and 10q-specific (*BlnI*) restriction sites in reads aligned to the 4q and 10q D4Z4 repeat arrays (**Fig. 1D**).

**Figure 1.**
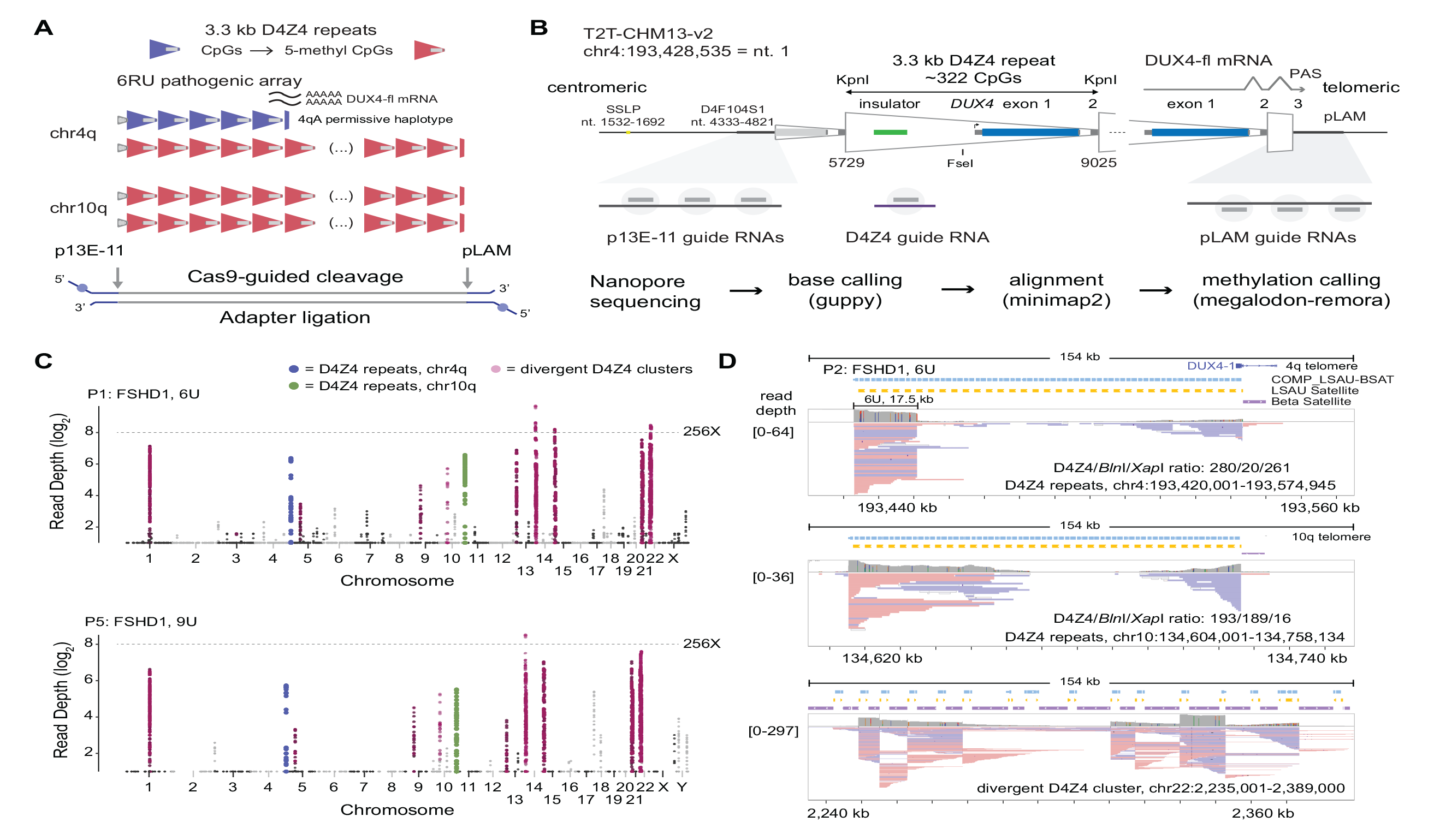
Cas9-targeted nanopore sequencing of D4Z4 repeats. **(A)** Schematic of D4Z4 repeat arrays found in sub-telomeric regions of human chr4q and 10q. A 6U D4Z4 array is shown on the permissive 4qA haplotype, and blue/red shading of repeats indicates CpG methylation. **(B)** Locations of Cas9-guide RNA cleavage sites at the centromeric (p13E-11) end, telomeric (pLAM) end, and within each repeat (D4Z4). Steps used in the sequencing pipeline for identifying targeted reads are shown. **(C)** Read depth Manhattan plots from two FSHD1 participants (P1, P5) mapped to the T2T CHM13 v2.0 reference genome (log_2_ scale, minimum depth = 2 reads, summarized in 2 kb, non-overlapping windows). Colors indicate reads mapped to the chr4q (blue) and 10q (green) D4Z4 arrays, as well as divergent *DUX4* clusters (purple). **(D)** IGV browser alignments of targeted reads from FSHD1 participant P5 with a 4qA 6U contraction. The D4Z4/*Bln*I/*Xap*I ratio shows the number of 3.3 kb D4Z4 repeats contained with the aligned reads and the number of 10q-specific (*Bln*I) and 4q-specific (*Xap*I) restriction sites found within those repeats. The lower panel shows alignments to a divergent D4Z4 repeat cluster on chr22.

**Table 1.** Read depth and targeting efficiency from nanopore Cas9 targeted sequencing in FSHD1 and FSHD2 participants and healthy controls

| Participant ID | Total Mapped Reads |  |  |  |  |  |  |  |  |  |  |  |  |  |
| --- | --- | --- | --- | --- | --- | --- | --- | --- | --- | --- | --- | --- | --- | --- |
|  |  | All On-target Reads |  |  | 4q On-target Reads |  |  | 10q On-target Reads |  |  | D4Z4 On-target Reads (3.3kb) |  | Divergent D4Z4 Reads |  |
|  |  | Total | % p13-pLAM | % D4Z4 | p13 | pLAM | Full Array | p13 | pLAM | Full Array | 4q | 10q | Reads | % Total |
| Cas9 gRNA:p13E-11+pLAM<sup>a</sup> |  |  |  |  |  |  |  |  |  |  |  |  |  |  |
| P1 | 35,019 | 392 | 1.1% | - | 91 | 78 | 68 (6U) | 109 | 114 | 38 (11U)<br>34 (13U) | - | - | 16,368 | 46.7% |
| P2 | 30,954 | 207 | 0.7% | - | 64 | 70 | 39 (6U) | 35 | 38 | 5 (13U) | - | - | 17,631 | 57.0% |
| P3 | 116,981 | 613 | 0.5% | - | 137 | 183 | 78 (6U) | 143 | 150 | 36 (9U) | - | - | 42,885 | 36.7% |
| P4 | 77,047 | 319 | 0.4% | - | 83 | 107 | 50 (6U) | 59 | 70 | 7 (13U)<br>1(19U) | - | - | 23,604 | 30.6% |
| P5 | 18,256 | 179 | 1.0% | - | 62 | 36 | 15 (9U-S)<br>10 (9U-L) | 56 | 25 | 0 | - | - | 8,381 | 45.9% |
| P6 | 27,317 | 139 | 0.5% | - | 48 | 54 | 36 (3U) | 14 | 23 | 1 (13U) | - | - | 9,044 | 33.1% |
| P7 | 53,538 | 385 | 0.7% | - | 30 | 34 | 4 (19U) | 171 | 150 | 63 (8U)<br>34 (12U) | - | - | 13,583 | 25.4% |
| P8 | 52,066 | 254 | 0.5% | - | 33 | 42 | 11 (16U)<br>1 (26U) | 87 | 92 | 39 (10U)<br>1 (25U) | - | - | 20,093 | 38.6% |
| Cas9 gRNA:p13E-11<sup>a</sup> |  |  |  |  |  |  |  |  |  |  |  |  |  |  |
| P2 | 30,734 | 77 | 0.3% | - | 27 | 0 | 0 | 50 | 0 | 0 | - | - | 42 | 0.1% |
| Cas9 gRNA:D4Z4<sup>b</sup> |  |  |  |  |  |  |  |  |  |  |  |  |  |  |
| C1 | 9,042 | 2,741 | - | 30.3% | - | - | - | - | - | - | 1,077 | 1,664 | 512 | 5.7% |
| C2 | 10,345 | 2,555 | - | 24.7% | - | - | - | - | - | - | 1,424 | 1,131 | 321 | 3.1% |
| P9 | 6,431 | 410 | - | 6.4% | - | - | - | - | - | - | 217 | 193 | 138 | 2.1% |
| P10 | 1,418 | 476 | - | 33.6% | - | - | - | - | - | - | 258 | 218 | 66 | 4.7% |
| P1 | 6,977 | 826 | - | 11.8% | - | - | - | - | - | - | 372 | 454 | 246 | 3.5% |
| P1 | 3,348 | 339 | - | 10.1% | - | - | - | - | - | - | 162 | 177 | 117 | 3.5% |
| Cas9 gRNA:p13E-11+pLAM+D4Z4 |  |  |  |  |  |  |  |  |  |  |  |  |  |  |
| P9<sup>a</sup> | 116,799 | 3,582 | 0.5% | 2.6% | 115 | 210 | 40 (6U) | 107 | 109 | 0 | 1,524 | 1,517 | 37,467 | 32.1% |
| P10<sup>a</sup> | 39,605 | 1,979 | 0.7% | 4.3% | 63 | 91 | 42 (6U) | 55 | 50 | 8 (13U) | 888 | 832 | 16,052 | 40.5% |
| C3<sup>c</sup> | 9,952 | 1,219 | 0.2% | 12.0% | 3 | 10 | 0 | 5 | 4 | 0 | 806 | 391 | 4,152 | 41.7% |
| Ultra -Long kit V14<sup>d</sup> |  |  |  |  |  |  |  |  |  |  |  |  |  |  |
| C4 | 6,140,482 | 253 | 0.004% | 0.037% | 74 | 51 | 14 (16U)<br>4 (41U) | 62 | 66 | 10 (20U)<br>5 (21U) | - | - | - | - |

<sup>a</sup>Sequencing on MinION R9.4.1 flowcell; <sup>b</sup> Sequencing on Flongle R9.4.1 flowcell; <sup>c</sup> Sequencing on MinION R10.4.1 flowcell; <sup>d</sup>Sequencing on PromethION 10.4.1 flowcell.

The additional enriched peaks outside of 4q and 10q telomeres (**Fig. 1C**) mapped to regions annotated in the T2T reference genome as LSAU-beta-satellite (BSAT) composite repeats that included homologous D4Z4 repeats (Hoyt et al. 2022). The location of these reads suggested targeting by the pLAM guide RNAs adjacent to divergent D4Z4 sequence, particularly in clusters of LSAU-BSAT repeats (**Fig. 1D**). The 20 largest genomic regions with LSAU-BSAT annotations (average span = 200 kb, range 20 kb – 621 kb, total span = 4.0 Mb; **Supplemental Table S3**) accounted for, on average, 39.2% of the total reads, ranging from 25.4 to 57.0% (**Table 1**, Divergent D4Z4 Reads). We confirmed that these reads were due to Cas9 cleavage at pLAM-related targets by removing the pLAM guide RNAs from a second library prepared from participant P2. This library retained the on-target p13E-11 reads but eliminated the LSAU-BSAT region reads (**Table 1**).

The high percentage of near-target reads at divergent D4Z4 arrays implied pLAM gRNAs were effectively cleaving multiple targets per genome equivalent. This suggested that a single gRNA designed within the D4Z4 repeat may yield a high percentage of monomer-sized, 3.3 kb D4Z4 reads arising from the 4q and 10q telomeres. We generated six additional libraries using a single D4Z4-gRNA (**Fig. 1B**) and observed a high yield of on-target, 3.3 kb monomer-sized, D4Z4 reads using the smaller Flongle flow cell (average = 19.5% on-target D4Z4 reads, **Table 1**). The ratio of monomer (3.3 kb) to dimer (6.6 kb) D4Z4 reads averaged 19:1, indicating almost complete cleavage of the D4Z4 arrays by the D4Z4 Cas9-gRNA complex. Three additional samples had p13E-11/pLAM and D4Z4 gRNA libraries prepared separately and combined for sequencing onto single MinION flow cells. These mixed libraries (p13E-11/pLAM and D4Z4) showed comparable yields for their component gRNA targets (**Table 1**). Finally, we compared the yield of targeted reads from the nCATs libraries sequenced on single MinION flow cells to a public dataset derived from a peripheral blood DNA library using ultra-long protocols and sequenced on three PromethION flow cells. This library had a total read yield 120X greater than our average MinION nCATs library, and a total Gigabase yield that was 450X greater (**Table 1**, sample C4). Despite this large difference in the overall yield of the PromethION dataset, comparable reads counts localized to the 4q and 10q D4Z4 arrays (**Table 1**) suggesting that the lower cost nCATs methodology is currently on par with whole genome nanopore sequencing for specifically studying the D4Z4 regions.

### D4Z4 arrays have asymmetric CpG methylation gradients

To evaluate the extent of hypomethylation that occurs as a result of contraction of the D4Z4 array, we extracted 5-methyl CpG (5mC) base modification calls from the nanopore reads. The methylation status of individual p13E-11 to pLAM cut-to-cut reads was summarized at the D4Z4 repeat level, using approximately 320 CpG dinucleotides per 3.3 kb repeat. The results were quantified as the percentage of 5mC per D4Z4 repeat and the distributions for 4qA and 10q arrays from FSHD1 participants P1 and P3 are shown as box plots in **Fig. 2A**. Repeat-level methylation increases in a step-wise fashion from the centromeric (p13E-11) to telomeric (pLAM) repeat for 6U, 9U, 11U, and 13U arrays. Smoothed methylation frequency plots and single-molecule methylation plots (**Fig. 2B**) using the same data displayed a characteristic striped methylation pattern within each repeat indicating a recurring pattern of higher and lower methylation, and a gradual, step-wise methylation increase across the arrays.

**Figure 2.**
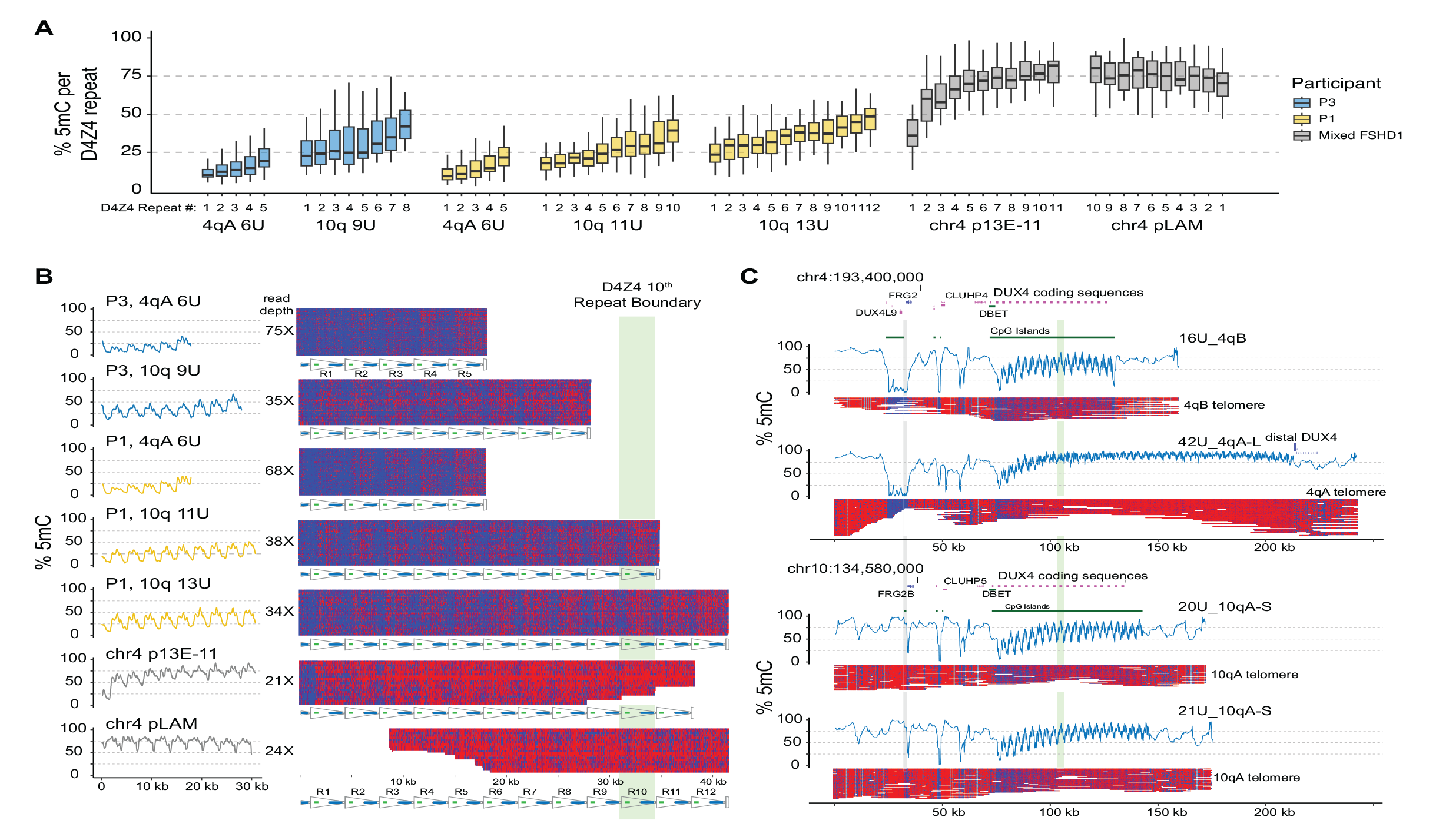
CpG methylation gradients traverse chr4q and 10q D4Z4 arrays. **(A)** Methylation levels of p13E-11 to pLAM targeted reads summarized as % 5mC sites per D4Z4 repeat (3.3 kb KpnI-to-KpnI region with ∼320 CpG sites per repeat). Repeat numbering begins at the centromeric end (p13E-11) and is reversed for the chr4 pLAM (telomeric) reads. **(B)** Single-read plots of targeted reads from (A) using *modbamtools* with unmethylated CpGs represented in blue and 5-methyl CpGs in red. The schematic representation of D4Z4 repeats shows *DUX4* exons 1 and 2 and the CTCF insulator region, and the green stripe indicates the 10^th^ centromeric D4Z4 repeat. (**C**) Alignments and methylation plots of untargeted reads (R10.4.1_e8.2 nanopores) from a control subject (C4) mapped to the 4qB 16U haplotype, the 4qA 43U haplotype, the 10q 20U haplotype, and the 10q 21U haplotype. The grey stripe delineates the start of the syntenic telomeric region shared between 4q and 10q, and the green stripe marks the 10^th^ repeat.

To examine the methylation levels of longer D4Z4 arrays in individuals with FSHD1, we aggregated the p13E-11 or pLAM reads from non-contracted 4qA alleles. The results showed that the centromeric end (p13E-11) had higher levels of methylation and a steeper increase than the shorter arrays, while the telomeric end (pLAM) had a uniformly high level of methylation with a slight gradual decrease distally (**Fig. 2A,B**). To ensure that these patterns were not specific to individuals with FSHD1, we also analyzed whole-genome data from a publicly available healthy control (C4) sequenced on high-density PromethION flow cells, where both alleles on chromosome 4 and 10 could be resolved. The whole genome nanopore data allowed us to assemble reads across the entire syntenic chromosome 4 and 10 region, and asymmetric methylation gradients were observed on all four haplotypes: the 4qB 16U, 4qA 42U, 10q 20U, and 10q 21U (**Fig. 2C**). The data confirmed the presence of an asymmetric methylation pattern in this control individual, indicating that these asymmetric methylation gradients are not specific to FSHD1.

### Methylation gradient modeling and methylation spreading

Despite the consistent gradient from low methylation in the proximal end of the array and higher methylation in the distal end, we observed a high level of variability in overall methylation between reads and within reads from one repeat to the next (**Fig. 3A**). To better capture variation in the methylation gradient, we used a linear mixed-effects model (LMM) based on read-level D4Z4 repeat unit methylation as the repeated measure. The LMM estimated the %5mC level from the first D4Z4 repeat (intercept) and the change per repeat (slope). For participant P1, the LMM estimated an intercept of 9.7% 5mC at the first D4Z4 unit with an increase in 3.7% 5mC per D4Z4 repeat (slope) using 68 full 4qA 6U reads (**Fig. 3A**). The LMM also allowed for the calculation of separate intercepts and slopes for individual reads, and these read-level regression lines are plotted with their measured repeat-level %5mC (**Fig. 3A**). The 4q and 10q arrays from FSHD1 participants with full-length reads (4q: 3U, 6U, 9U and 10q: 9U, 11U, 13U) had intercepts from 9.6 to 26.1% and slopes from 1.9 to 3.9% (**Table 2**, **Fig. 3B**). FSHD2 participants (4q:16U, 19U and 10q 8U, 10U, 12U) showed intercepts ranging from 9.1 to 14.3%, similar to FSHD1 participants, but showed lower slopes, ranging from 0.7 to 1.6% (**Table 2**, **Fig. 3B**).

**Figure 3.**
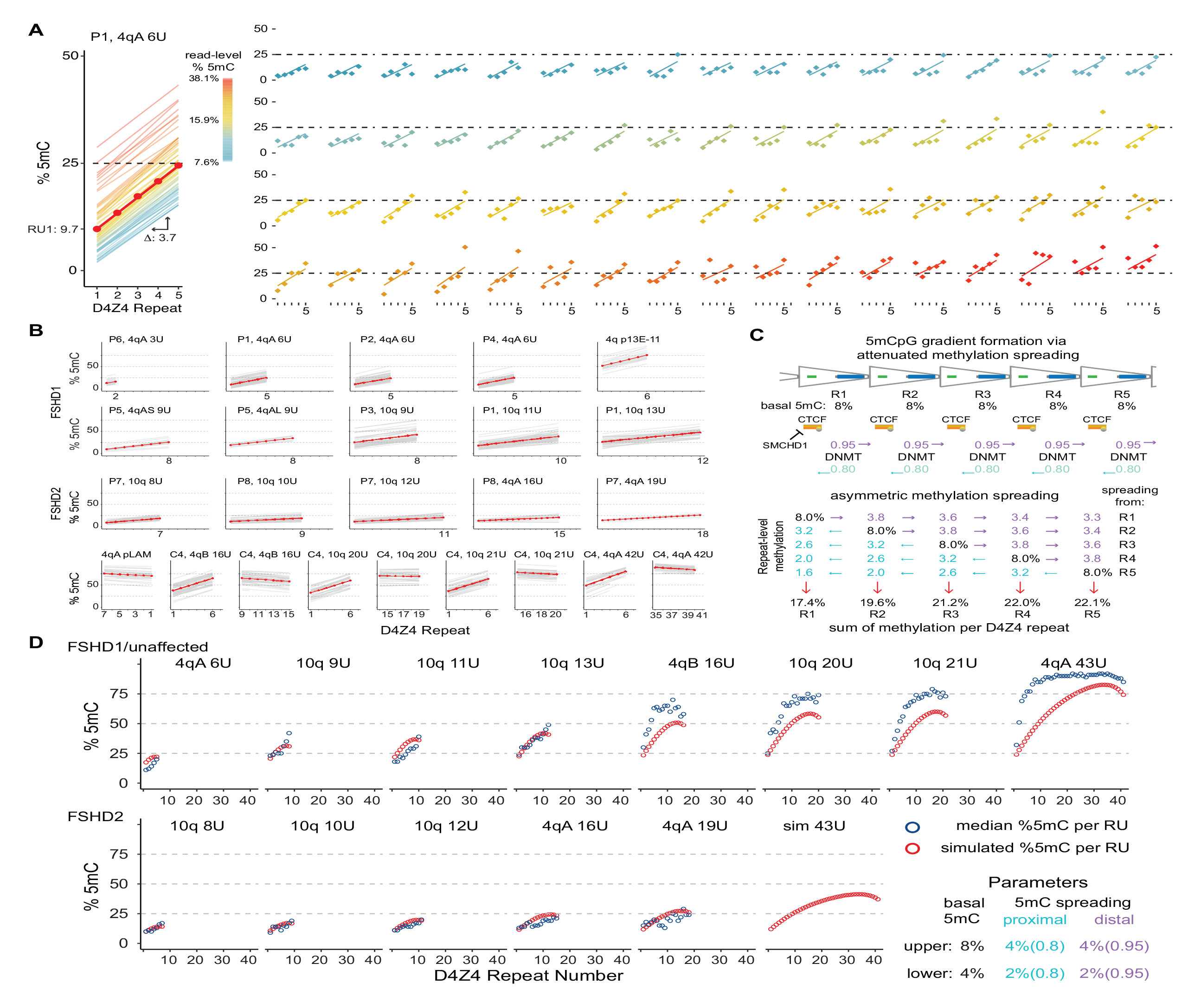
Methylation gradient modeling. (**A**) Intercept and slope estimates from p13E-11 to pLAM 4qA 6U reads from participant P1. The red line is the model estimate and the thin lines are the individual read estimates with a color scale for read-level total % 5mC. The right panel shows the observed D4Z4 methylation frequencies for each read (points) with their individual read estimates. The 68 plots are ordered by read-level % 5mC. (**B**) Intercept and slope estimates as in (A) from FSHD1, FSHD2 participants, and control subject C4 (see Table 2). (**C**) Model for gradient formation via basal methylation followed by asymmetric spreading. (**D**) Predicted D4Z4 methylation levels from simulated basal methylation and spreading (red points) compared to observed values (blue points, mean methylation per repeat, data from (B)). In the lower panel, the basal methylation and spreading parameters were reduced by a factor of 2.

**Table 2:** D4Z4 methylation gradient analysis using a linear mixed-effects model to estimate intercept and slope.

| Subject ID | Chromosome | Length D4Z4 | % Methylated CpG RU1 |  | Change in % methylated CpG per RU<sup>a</sup> |  |
| --- | --- | --- | --- | --- | --- | --- |
|  |  |  | Intercept | 90% CI | Slope | 90% CI |
| FSHD1 |  |  |  |  |  |  |
| P6 | chr4qA-S | 3U | 14.2 | 10.6 - 17.7 | 0 | -2.1 - 2.1 |
| P1 | chr4qA-S | 6U | 10.3 | 7.3 - 13.2 | 2.8 | 2.3 - 3.4 |
| P2 | chr4qA-S | 6U | 15.3 | 12.0 - 18.5 | 2.1 | 1.5 - 2.8 |
| P3 | chr4qA-S | 6U | 11.5 | 8.6 - 14.5 | 2.3 | 1.8 - 2.8 |
| P4 | chr4qA-S | 6U | 10.4 | 7.2 - 13.7 | 2.9 | 2.3 - 3.5 |
| P5 | chr4qA-S | 9U | 10.1 | 5.6 - 14.5 | 2 | 1.3 - 2.7 |
| P5 | chr4qA-L | 9U | 18 | 12.8 - 23.2 | 2.3 | 1.4 - 3.1 |
| P3 | chr10q | 9U | 24.9 | 21.5 - 28.3 | 2.3 | 1.7 - 2.8 |
| P1 | chr10q | 11U | 17.4 | 15.1 - 19.7 | 2.2 | 1.9 - 2.6 |
| P1 | chr10q | 13U | 25.4 | 22.0 - 28.8 | 1.9 | 1.3 - 2.4 |
| P1, P2, P3, P4, P5, P6 | chr4qA-S | >10U, centromeric | 46.6 | 42.5 - 50.7 | 6.1 | 5.3 - 6.8 |
| P1, P2, P3, P4, P5, P6 | chr4qA-S | >10U, telomeric | 76.2 | 72.1 - 80.3 | -1 | -1.6 - -0.3 |
| FSHD2 |  |  |  |  |  |  |
| P7 | chr10q | 8U | 9.5 | 6.5 - 12.5 | 1.2 | 0.7 - 1.7 |
| P8 | chr10q | 10U | 11.9 | 8.6 - 15.2 | 0.7 | 0.2 - 1.3 |
| P7 | chr10q | 12U | 11.8 | 8.4 - 15.3 | 0.8 | 0.2 - 1.3 |
| P8 | chr4qA-S | 16U | 13.3 | 8.4 - 18.2 | 0.5 | -0.3 - 1.2 |
| P7 | chr4qA-S | 19U | 14.4 | 7.0 - 21.8 | 0.6 | -0.5 - 1.7 |
| Unaffected Control |  |  |  |  |  |  |
| C4 | chr4qB | 16U, prox 6U | 33.7 | 30.4 - 37.0 | 6.2 | 5.6 - 6.8 |
| C4 | chr4qB | 16U, distal 6U | 64.1 | 60.7 - 67.4 | -1.3 | -1.9 - -0.7 |
| C4 | chr10q | 20U, prox 6U | 29.4 | 25.9 - 33.0 | 5.7 | 5.1 - 6.4 |
| C4 | chr10q | 20U, distal 6U | 71.4 | 67.9 - 74.9 | -0.5 | -1.2 - 0.1 |
| C4 | chr10q | 21U, prox 6U | 34.5 | 31.0 - 37.9 | 5.7 | 5.1 - 6.4 |
| C4 | chr10q | 21U, distal 6U | 77.1 | 73.5 - 80.6 | -0.9 | -1.6 - -0.2 |
| C4 | chr4qA-L | 42U, prox 6U | 42.5 | 38.4 - 46.6 | 7.1 | 6.3 - 7.9 |
| C4 | chr4qA-L | 42U, distal 6U | 89.8 | 86.3 - 93.4 | -1 | -1.6 - -0.4 |

<sup>a</sup>RU-repeat unit

Variation in the overall per read methylation was evident, as well as a trend for increased methylation of the first D4Z4 repeat with longer arrays (**Fig. 3B**, **Table 2**). Centromeric (p13E- 11) reads from longer arrays had the highest intercepts, including the >10U, 4q reads from FSHD1 participants (52%) and the 16U to 42U arrays from the control subject, which ranged from 32.7 to 48.5%. Telomeric reads (pLAM) from the longer arrays consistently showed negative slopes, ranging from −1.1 to −0.2, suggesting a tapering from the high %5mC levels seen for internal repeats in very large arrays.

These patterns suggested a simple model combining basal D4Z4 methylation and spreading that could lead to the formation of asymmetric gradients. **Fig. 3C** shows a model based on *ad hoc* parameters for basal methylation rates (8% per D4Z4 repeat) and asymmetric spreading (a directional rate of 95% distal spreading versus 80% proximal spreading) that results in asymmetric gradient formation. We used these parameters to simulate D4Z4 methylation levels for different length arrays. The modeled findings were consistent with the observed median D4Z4 repeat levels from FSHD1 and control participants, including the tapering of methylation in the longer arrays (**Fig. 3D**). When the basal parameter was reduced by a factor of 2, from 8% to 4%, the simulated gradient more closely paralleled the observed methylation levels in FSHD2 participants.

### Fine-scale D4Z4 methylation patterns suggest nucleosome positioning

In addition to methylation gradients across the D4Z4 array, we observed consistent methylation patterns within individual repeats, including areas of low and high methylation. The single-read and smoothed methylation frequency plots of the 4qA 6U haplotype reads from participant P1 (**Fig. 4A**) demonstrate this repeating pattern across the array from R1 to R5. Aligning methylation frequencies of individual CpGs within the 6U haplotype from participant P1 with the annotated features of the 3.3 kb *Kpn*I-*Kpn*I repeat showed recurring low and high methylation levels across repeats R1-R5 (**Fig. 4B**). The smoothed methylation frequency from repeats R1 to R5 showed a consistently rising pattern of methylation with repeat number. The CTCF/insulator region were among the least methylated CpG residues, and this region contains the DR1 amplicon previously identified as relatively hypomethylated (Hartweck et al. 2013; Roche et al. 2019). CpGs near the *DUX4* transcription start site, especially the CpG at nucleotide (nt) 1704, had high methylation levels ranging from 28-90%. A phasic methylation pattern with ∼180 nt periodicity started near this region and propagated through *DUX4* exons 1 and 2 (**Fig. 4B**).

**Figure 4.**
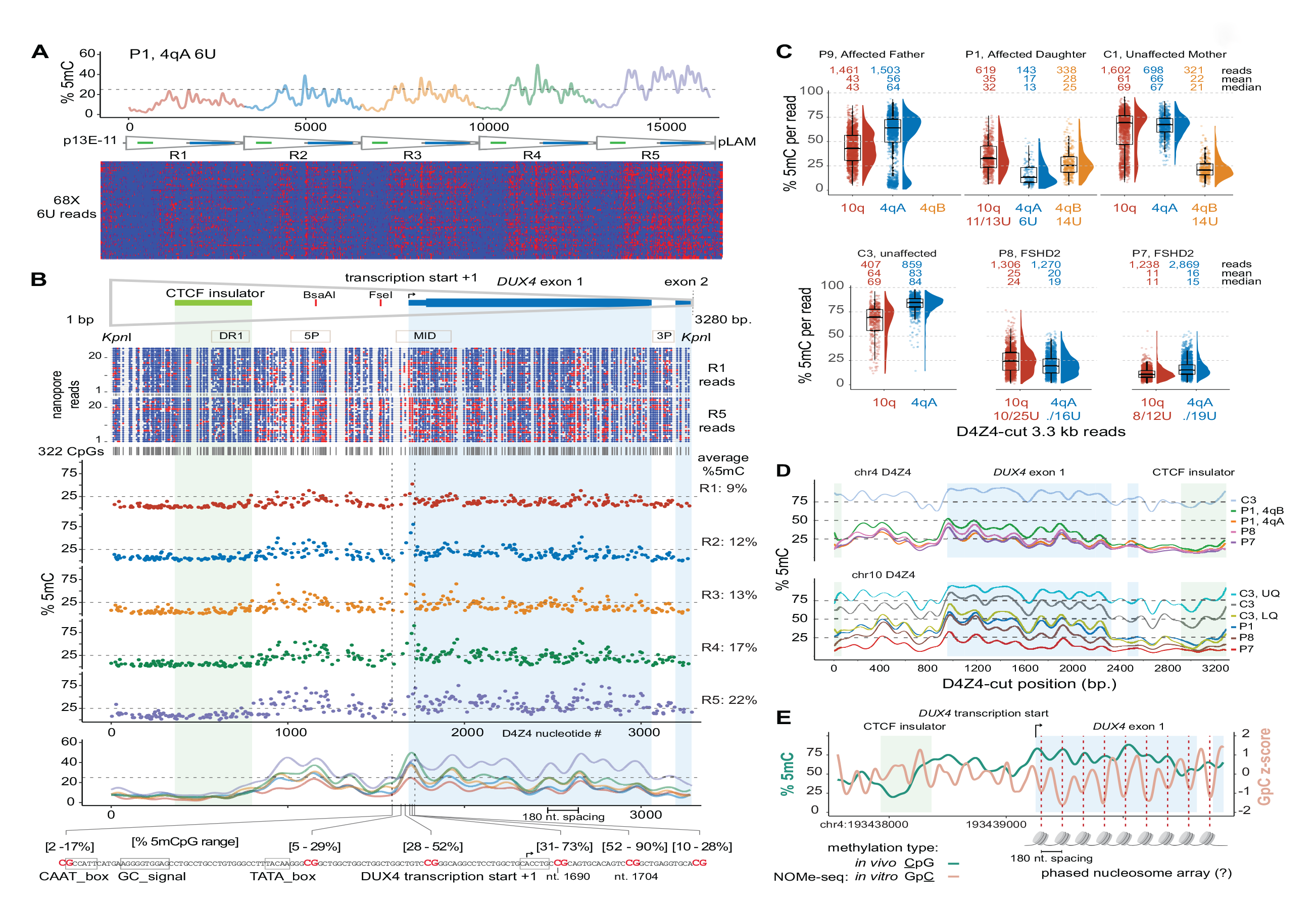
Fine-scale D4Z4 repeat methylation patterns. **(A)** Methylation plot of 4qA 6U reads from participant P1 using the “smooth_ksmooth” function (smoothness = 10) with single-read plots from *modbamtools*. **(B)** Annotated D4Z4 features including *DUX4* exons 1 and 2 (mRNA from AF117653.3), the CTCF insulator (*Blp*1-*Blp*1 fragment from Ottaviani *et al*.,2010), *BsaA*I and *Fse*I methylation-sensitive restriction sites, and PCR amplicons commonly used for bisulfite sequencing. A *methylartist* plot shows 322 CpG residues represented as blue (unmethylated) or red (methylated) dots for the R1 and R5 repeats from a subset of reads in (A). The middle panel shows the average individual CpG methylation from the 4qA 6U reads in (A) for the 322 CpGs, aligned from R1 to R5. The lower panel shows the methylation frequency plot smoothed as in (A), with the methylation range of six CpGs in the *DUX4* proximal promoter with the transcription start from Dixit *et al*. (AF117653.3). **(C)** Methylation levels of individual 3.3 kb D4Z4-cut reads with % 5mC per read on the y-axis. The number of reads and mean/median % 5mC per read are shown for each chromosome. Upper plot = FSHD1 trio (affected father, affected daughter, unaffected mother) segregating a 6U repeat; lower plot = unaffected versus FSHD2 participants. **(D)** Methylation frequency plots of D4Z4-cut reads from (C), smoothed as in (A). **(E)** % 5mC and Nucleosome Occupancy and Methylation sequencing (NOMe-seq) GpC z-scores plotted for the second chr4 D4Z4 repeat (centromeric) using *in vivo*/*in vitro* methylation data from the HG002 lymphoblastoid cell line.

D4Z4 3.3 kb reads from an enrichment using the Cas9-gRNA targeting individual D4Z4 repeats (**Fig. 1B**, **Table 1**) were used to examine the fine-scale methylation structure in additional FSHD1, FSHD2, and control participants. Although these reads lacked positional information for the individual D4Z4 repeat, sequence alignments enabled discrimination between 4qA, 4qB, and 10q repeats. Paternally inherited 4qA reads from the 6U haplotype carried by participant P1 were distinguishable from the 4qB reads inherited from her mother (**Fig. 4C**). The lower methylation range of these 4qA-specific reads and the intermediate methylation range of 10q reads from her 11U/13U haplotypes were similar to the repeat-level summaries from full haplotype reads (**Fig. 4B,C; Fig. 3B**). The 6U haplotype methylation status in her father was evident from the bi-phasic profiles of his 4qA-specific reads, emphasizing the distinction in repeat-level methylation between small and large arrays. The relative absence of higher methylation levels from the two FSHD2 participants was apparent, although the mean and median methylation levels of the FSHD2 4qA reads were comparable to the 4qA reads from the 6U haplotype from FSHD1 participant P1. The smoothed methylation frequency plots of the 4qA D4Z4 reads showed that the methylation profiles of FSHD1 and FSHD2 were similar (**Fig. 4D**). This result suggests that reductions overall methylation levels specific for FSHD2 does not alter the fine-scale D4Z4 methylation pattern. From both 4q and 10q profiles, the maintenance of this fine-scale pattern with increasing methylation is evident, and the phasic methylation pattern beginning near the start of *DUX4* exon 1 was seen with repeats from both chromosomes and with increasing overall methylation levels.

The 180-nt periodicity of CpG methylation suggested the possibility that this phasic pattern corresponds to nucleosome occupancy. To test this hypothesis, we examined the D4Z4 region for nucleosome protection utilizing a nanopore NOMe-seq data set (Nucleosome Occupancy and Methylome Sequencing) generated from HG002 lymphoblastoid cells by the T2T consortium (Gershman et al. 2022). The CpG methylation frequency in the extended 4qA region of the HG002 cell line was comparable to the 4qA 42U allele from peripheral blood of control sample C4 and an asymmetric gradient was seen at the centromeric end of the HG002 array (**Supplemental Fig. S1**). This methylation gradient was also observed in the profile from HG002 PacBio HiFi long-read sequencing, confirming that these methylation gradients can be detected with an orthogonal long-read sequencing chemistry. The smoothed frequency plot from the 4q centromeric D4Z4 repeat R2 displayed an intermediate level of methylation and a smoothed frequency plot displayed a phasic profile similar to the profiles observed in peripheral blood (green line, **Fig. 4E**). The NOMe-seq GpC z-score, which measures the relative protection of GpC dinucleotides from *in vitro* methylation of isolated chromatin with a GpC methyltransferase (Kelly et al. 2012), also showed a ∼180 nt GpC phasic pattern across the exon 1 and 2 region (orange line, **Fig. 4E**). The GpC z-score valleys were coincident with CpG methylation peaks indicating that CpG methylation is higher in more protected regions. The endogenous phasic CpG methylation across exon 1 was offset by ∼90 nt from the GpC z-score, consistent with the methylation status of CpGs in this region influencing the positioning of nucleosomes.

### Divergent D4Z4 repeats including a paralogous chr22 cluster are highly methylated

The abundance of pLAM-related reads, as shown in Table 1, facilitated a genome-wide analysis of the sequence and methylation status of D4Z4-related sequences. We examined the copy number in the T2T reference genome of a 4 kb query sequence encompassing the distal 3.3 kb D4Z4 repeat with the terminal ∼0.3 kb D4Z4 truncation and the flanking ∼0.4 kb pLAM sequence, including the polymorphic PAS region permissive for DUX4 expression from the distal D4Z4 repeat (Lemmers et al. 2010a). The T2T reference genome contained 13 occurrences of this full 4 kb sequence, including the two copies on 4q and 10q. A ∼3.1 kb sequence that deleted ∼0.9 kb beginning at the 5’ *Kpn*I site including the CTCF/insulator region, was the most common D4Z4-related sequence with 319 copies (**Fig. 5A**). Both the full and 5’ truncated copies were primarily located in the LSAU-BSAT rich regions of acrocentric chromosomes (chr13, 14, 15, 21, and 22) and in the pericentric region of chr1 (**Supplemental Table S4**) and accounted for approximately 98% of the full-length, divergent *DUX4* ORFs in the T2T reference genome (**Supplemental Table S5**). A high methylation state of a full D4Z4- pLAM sequence from chr22 is shown in **Fig. 5B**, where a percent-identity plot indicates the ∼90% identity with the D4Z4-4qAS-pLAM region.

**Figure 5.**
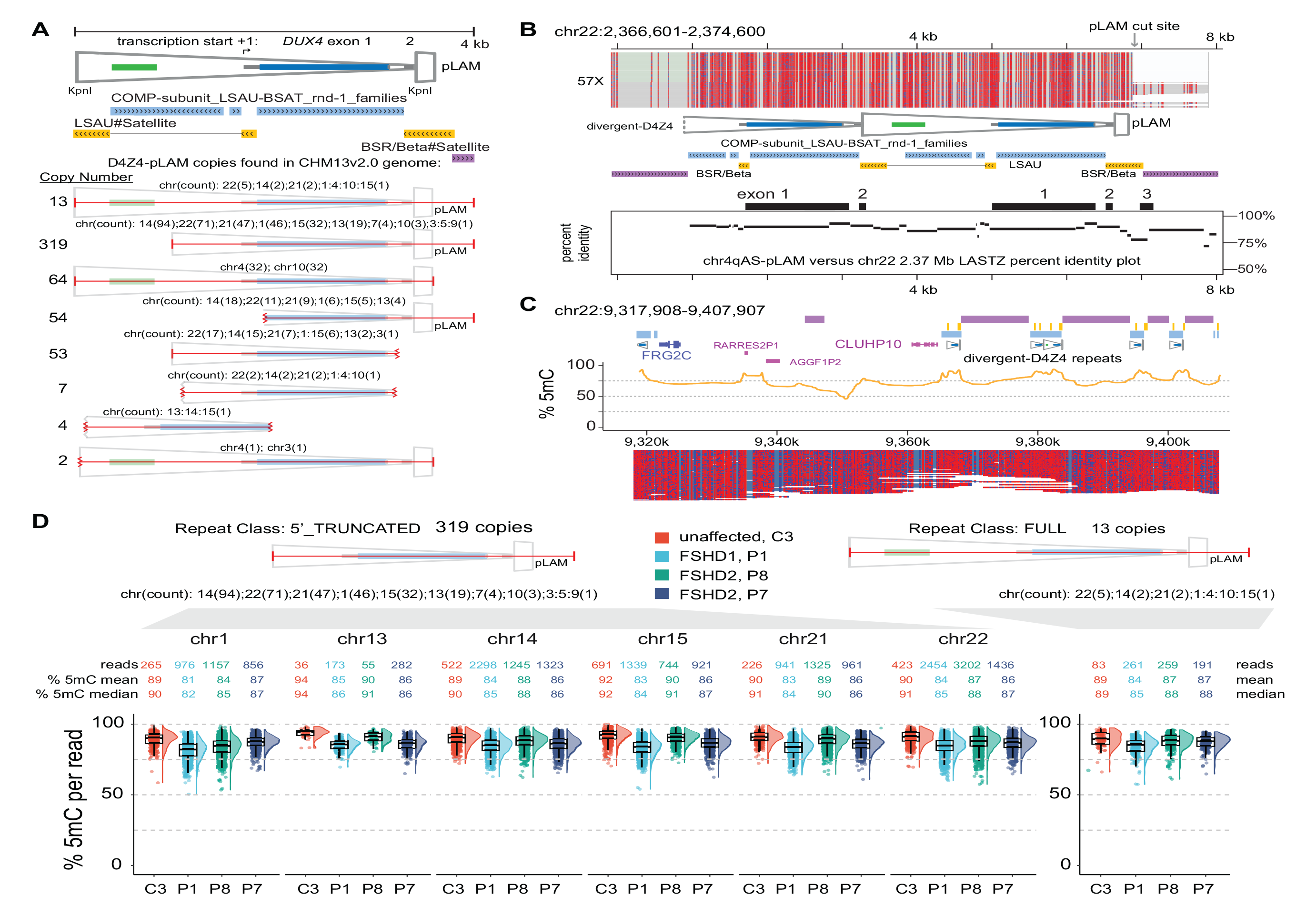
Methylation profiles of divergent D4Z4 repeats. **(A)** Annotated structure and copy number of the distal 4 kb D4Z4 repeat beginning at the 5’ *Kpn*I site through the 4qAS pLAM sequence/β-satellite junction (see Supplemental Table S2 for list of genomic locations). **(B)** Methylation profile of a full-length, divergent D4Z4 repeat found on chr 22. The LASTZ percent identity (PIP) plot shows the distribution of sequence identity along the length of this chr22 D4Z4 repeat compared to the distal chr 4qAS D4Z4 repeat. (**C**) Methylation plots of untargeted reads mapping to chr22:9,317,908-9,717,807 region from the C4 control (R10.4.1_e8.2 nanopores). **(D)** Distributions of CpG methylation levels of divergent D4Z4 repeats where each point represents the % 5mC of an individual read within the region that aligns completely with a divergent repeat. The C3 sample was sequenced using R10.4.1_e8.2 nanopores.

The T2T reference genome resolved numerous regions that contain segmental duplications of FSHD region gene 1 (*FRG1*) and FSHD region gene 2 (*FRG2*) (Nurk et al. 2022). We examined a T2T chr22 region with a divergent D4Z4 cluster that retains paralogous sequence through the *FRG2* gene and the inverted single D4Z4 repeat found on 4q but absent from 10q. The methylation plots reveal a highly methylated state for both full and 5’ truncated D4Z4 repeats in this region (**Fig. 5C**). A dot plot alignment of the 4q region and the entire ∼380 kb chr22 D4Z4 cluster demonstrates high sequence conservation that extends beyond the *FRG2* paralog (**Supplemental Fig. S2**). The divergent D4Z4 repeats in this cluster are interspersed with BSAT repeats and can be found in both forward and reverse orientations relative to 4q D4Z4 repeats. Moreover, the methylation frequency and single-read plots of the entire ∼380 kb chr22 D4Z4 cluster support a uniformly high level of methylation without evidence for methylation gradients (**Supplemental Fig. S2**).

In contrast to arrays on 4q and 10q, methylation levels for these divergent D4Z4 arrays were uniformly high. We conducted an analysis of methylation levels derived from reads aligned to the full and 5’ truncated class of divergent repeats across multiple chromosomal regions for FSHD1 participant P1, FSHD2 participants P7 and P8, and control participant C3. A total of 24,645 read alignments (803 full and 23,851 5’ truncated) were utilized and the median methylation levels for all four participants ranged from 81% to 94% (**Fig. 5D**). Notably, there were no read alignments with a methylation level below 50%, and only 11 reads demonstrated methylation levels below 60%. This observation indicates that the methylation levels of divergent D4Z4 repeats were uniformly high in the peripheral blood DNA of both FSHD1 and FSHD2 affected participants, as well as unaffected individuals.

### Methylation gradients are seen in other high CpG ‘continents’

To identify whether the methylation gradients identified in the 4q and 10q D4Z4 arrays were unique, we conducted a genome-wide search for other regions of comparable CpG density to assess whether these areas harbored similar methylation gradients. A 33 kb sliding window, equivalent in size to 10 D4Z4 repeat units, was scanned across the T2T reference genome using a step size of 3.3 kb (∼1 repeat unit) to rank the genome-wide CpG densities in 33 kb segments. CpG density for each individual 33kb segment was plotted against the location of the segment in a Manhattan plot with segments showing >7% CpG density highlighted for further analysis (**Fig. 6A**). The 4q and 10q D4Z4 arrays were the two most CpG-dense regions across the genome for this window length, with densities of approximately 9.8% for the 33 kb window. There were 8 additional regions with 33 kb windows of >7.0% CpG density (**Supplemental Table S6**). Five peaks were from 45S ribosomal DNA (rDNA) array clusters, and unique peaks were seen for the chr1 LM-tRNA cluster, the chr1 5S rDNA cluster, and the chr19 SST1 cluster. Methylation plots from the HG002 cell line showed evidence of methylation gradients in the rDNA arrays, the chr1 tRNA cluster, and the chr1 5S rDNA cluster (**Supplemental Fig. S3**).

**Figure 6.**
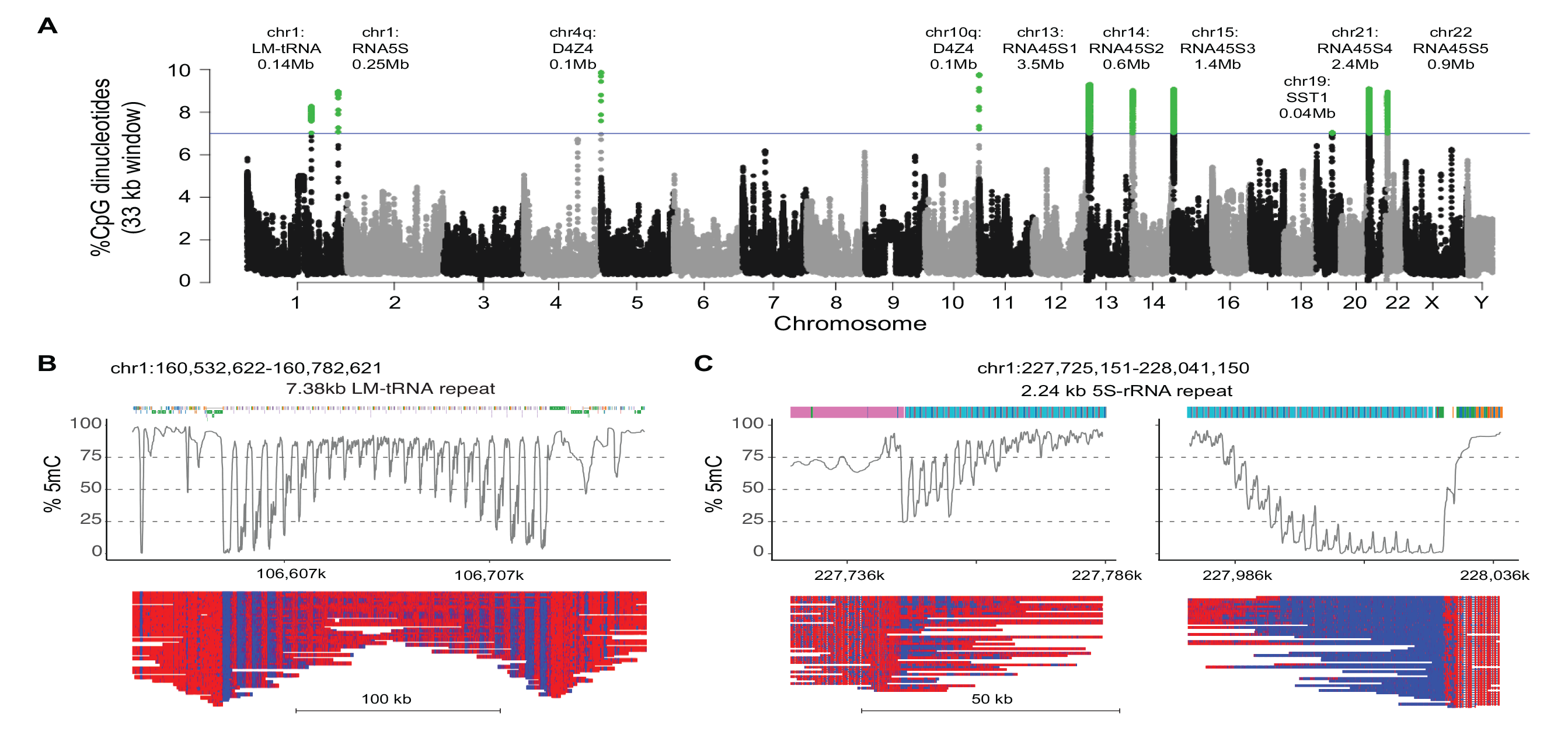
Methylation gradients in other CpG-dense repeat regions. **(A)** Manhattan plot of CpG density in 33 kb sliding windows (step size = 3.3 kb) across the T2T CHM13 v2.0 reference genome. Windows above 7% CpG density are marked as green dots. Alignment and methylation plots of untargeted reads from the C4 control (R10.4.1_e8.2 nanopores) mapped to **(B)** the tRNA cluster on chromosome 1 and **(C)** the 5S rRNA cluster on chromosome 1. Mapped reads are anchored by unique flanking sequence and the two ends of the 5S rRNA cluster are shown separately. The annotations are the RepeatMasker repetitive element UCSC browser track.

Since long-range assembly across the 45kb rRNA arrays remains challenging, we focused on assembly and methylation analysis from the chr1 tRNA and 5S rDNA cluster using the peripheral blood reads from control C4. The chr1 tRNA has a 7.4 kb LM-tRNA repeat and the 5S region has a 2.2 kb 5S-rRNA repeat, so we used only reads anchored uniquely into flanking sequence for methylation analysis. The LM-tRNA region had a symmetrical gradient on both ends of the repeat array (**Fig. 6B**), while the 5S-rRNA region had asymmetrical gradients (**Fig. 6C**), confirming the presence of methylation gradients in repeat segments outside of the 4q and 10q D4Z4 arrays. Notably, the chr1 LM-tRNA and 5S rRNA clusters were previously identified as hypomethylated regions in FSHD2 patients (Mason et al. 2017).

## DISCUSSION

FSHD is one of the most common muscular dystrophies, yet genetic analysis is among the most difficult of any monogenetic disorder, due to the complex mechanism combining genetic and epigenetic factors with long stretches of repetitive DNA that have close paralogs elsewhere in the genome. Southern blot techniques have been the gold standard for diagnosis and research analysis for many years, but recent development of optical mapping, molecular combing, and next-generation based bisulfite sequencing techniques are rapidly enhancing our understanding of the genetics of FSHD (Vasale et al. 2015; Nguyen et al. 2017; Roche et al. 2019; Zampatti et al. 2019; Dai et al. 2020; Gould et al. 2021).

Recently developed long read sequencing techniques such as nanopore sequencing, have opened the door to analysis of highly repetitive DNA in areas of the genome that were previously unmappable. Using these techniques, researchers completed sequencing of a full reference human genome for the first time, including mapping of repetitive DNA such as the D4Z4 arrays important in FSHD (Nurk et al. 2022). Initial efforts to use nanopore sequencing for FSHD, struggled due to lack of read-depth in 4q derived D4Z4 arrays from genomic analyses (Mitsuhashi et al. 2017). Development of strategies to enrich sequences of interest using Cas9-targeting has led to improved read-depth for regions of interest (Gilpatrick et al. 2020), including the D4Z4 array associated with FSHD (Hiramuki et al. 2022) Nanopore sequencing not only opens the door to the full sequencing of the D4Z4 array, but also allows determination of CpG methylation in the array, a critical aspect of genetic analysis for FSHD.

Here we described a refinement of the Cas9 targeting technique that can successfully enrich 4q and 10q D4Z4 arrays and provide a comprehensive tool for genetic analysis of FSHD. We demonstrated on-target p13E-11-to-pLAM reads enriched to 0.5-1.1% of all mapped reads, which provides sufficient read depth for analysis of 4q versus 10q sequences, determination of A/B haplotype, and size and methylation status of the array. Surprisingly, the bulk of off-target reads were near-target reads that arise from divergent D4Z4 arrays scattered throughout the genome, accounting for 25-57% of total reads. Exclusion of pLAM guide RNAs from the assay removed nearly all of these near-target reads, but did not increase recovery of full-length reads. The high read-depth of near-target sequences from pLAM gRNAs suggested high efficiency Cas9 cleavage per genome equivalent.

Our investigation revealed that the use of a single gRNA within the D4Z4 repeat led to a substantial retrieval of D4Z4 repeat reads, with up to 30% on-target reads, >95% of which were 3.3 kb monomer, single D4Z4 reads. These findings offer additional support for the high efficiency of Cas9 cleavage. While the order of the individual repeats is lost using D4Z4 targeted guide RNAs, this assay adds read-depth and maintains the ability to separate 4qA, 4qB, and 10q reads, and quantify methylation. This assay can be done on lower cost Flongle flow cells, providing a quick screen to differentiate FSHD and healthy controls. Furthermore, the retrieval of uniform 3.3 kb reads offers a semi-quantitative measure of D4Z4 copy number, as their yield is not subject to length-dependent bias commonly seen in nanopore sequencing. A combined approach using libraries with p13E-11 and pLAM guide RNAs and libraries with D4Z4 guide RNAs on the same MinION flowcell provided adequate read-depth for comprehensive analysis and maintained the advantages of both assays. The combined assay provided read-depth comparable to non-targeted reads obtained from whole genome sequencing in a healthy control based on multiple PromethION flow cells obtained at a substantially higher cost.

The ability of nanopore sequencing to provide the methylation status of individual CpG sites on individual molecules allowed us to observe asymmetric methylation gradients that form at the proximal end of D4Z4 repeat arrays and reach a saturation point at approximately the 10^th^ repeat. This finding provides a biophysical correlate for why contractions of less than 10 repeats are associated with FSHD. We postulated a simple model for directional barriers to methylation spreading across the 4q and 10q arrays that would establish the observed length-dependent, asymmetric gradients. While *ad hoc*, the model predicts both the presence of asymmetric methylation gradients and the length-dependent change. The absence of methylation gradients and the uniform hypermethylation observed in paralogous D4Z4 clusters may provide insight to elements that restrict or promote methylation spreading.

It has been shown that methylation at the single *Fse*I site in the proximal (centromeric) D4Z4 repeat is correlated with repeat array size (Lemmers et al. 2015). The authors proposed a metric (Delta1 score) based on proximal *Fse*I methylation and a non-linear relationship between the cumulative array size, and used this as a marker for disease onset and progression. The length-dependent effect modeled by the Delta1 score was confirmed in our study by the use of a linear mixed-effect model to estimate the degree of change in the intercept value of the methylation gradient with increasing array size. The Delta1 score is particularly sensitive to detecting hypomethylation due to *SMCHD1* mutations in FSHD2 individuals due to hypomethylation of both 4q and 10q alleles. The hypomethylation effect on all 4q and 10q repeats of *SMCHD1* mutations seen with single D4Z4 guide RNA libraries and the decreased slope of the methylation gradient we observed in FSHD2 individuals provides an alternative method for quantifying D4Z4 methylation. Although fitting the intercept and slope using a linear mixed-effect model allowed for the quantification of haplotype-specific methylation levels, this approach only accounted for a part of the considerable variability in read-level methylation observed with different haplotypes. Alternative approaches to quantify this read-to-read variation may be useful as metrics for association with clinical markers of disease.

Bisulfite sequencing and methylated DNA immunoprecipitation analysis have been widely used for high resolution methyl-CpG analysis of the D4Z4 repeat (de Greef et al. 2009; Hartweck et al. 2013; Gaillard et al. 2014; Jones et al. 2014). These studies have been recently refined with improvements in amplicon specificity for discriminating 4q and 10q alleles, and by NGS-based analysis of bisulfite converted amplicons (Huichalaf et al. 2014; Roche et al. 2019; Gould et al. 2021). The methyl-CpG patterns observed in our study confirmed several aspects of these prior studies, including extended hypomethylation at the 5’ end of the D4Z4 repeat including the CTCF/insulator region and the DR1 region. In a new finding, we identified preferential methylation of certain CpGs in close proximity to the *DUX4* transcription start site and a consistent pattern of methylation across all ∼320 individual CpGs within each 3.3 kb repeat. This recurrent methylation pattern was independent of 4q or 10q haplotypes, FSHD1, FSHD2, or unaffected status, and was evident in both averaged single-CpG methylation estimates and kernel-smoothing analysis. Several aspects of this fine-scale methylation pattern suggested that it may be related to the chromatin state of the D4Z4 repeat. Two regions of relative hypomethylation were consistently observed at positions 100 to 700, overlapping with the CTCF/insulator region (Ottaviani et al. 2009), and from 1400 to 1700, corresponding to the D4Z4 binding element (DBE) site for YY1/HMGB2/NCL complexes situated in the *DUX4* proximal promoter region (Gabellini et al. 2002). Recent affinity capture of protein complexes bound to D4Z4 repeats suggest that these regions may also participate in binding the Nucleosome Remodeling Deacetylase (NuRD) and Chromatin Assembly Factor 1 (CAF-1) complexes (Campbell et al. 2018).

A phasic methylation pattern with a periodicity of approximately 180 nts was observed starting at the preferentially methylated CpGs near position 1700. This pattern was also detected using long-read sequencing data from the HG002 lymphoblastoid cell line, which included companion NOMe-seq data to concurrently investigate nucleosome positioning and CpG methylation. NOMe-seq data from a proximal D4Z4 repeat with an intermediate level of CpG methylation suggested that the phasic CpG methylation signal was consistent with a phased nucleosome array in which the more highly methylated CpG peaks were wrapped around nucleosomes. Although it is unclear how phased nucleosomes may contribute to *DUX4* repression, we can speculate that the preferentially methylated CpGs near position 1700 may participate in positioning of nucleosomes by localizing NuRD complexes through their methyl-CpG-binding protein domain 2 (MBD2) subunits to specific methyl-CpGs. Recent studies indicate that MBD2 has high affinity for large and densely methylated CpG islands, allowing recruitment of NuRD to facilitate nucleosome positioning, histone deacetylation, and chromatin compaction (Leighton et al. 2022). Further investigation is required to determine whether the fine-scale methylation pattern observed within each D4Z4 repeat is established during development or whether it arises from the dynamic turnover of methyl-CpGs, reflecting changes in the underlying chromatin state.

The T2T reference genome enabled us to map near-target reads from the pLAM gRNAs to the repetitive regions of acrocentric chromosomes and analyze the methylation state of paralogous D4Z4 copies. Our findings were consistent with previous studies, which suggest that the D4Z4-4qAS-pLAM sequence is the ancestral form of the D4Z4 retroelement that radiated and diverged during primate and human evolution (Lemmers et al. 2010b). The paralogous D4Z4 sequences we catalogued in the T2T genome uniformly share ∼90% nucleotide identity with the D4Z4-4qAS-pLAM sequence but with reduced CpG densities of approximately 5.4%. Notably, all these paralogous sequences showed high levels of methylation, regardless of length or location in the genome, and irrespective of whether the individual had FSHD1, FSHD2, or was unaffected. This hypermethylation was also observed in a large 380 kb cluster on chromosome 22, which has features similar to the 4q D4Z4 region, including an inverted *DUX4c* paralog located before a *FRG2* paralog at one end, and a full-length D4Z4-4qAS-pLAM paralog at the other end.

One of the more striking aspects of the 4q D4Z4 array is its extreme CpG density. We used this property to scan the genome with a window size of 33 kb for other large regions of extreme density. The 4q and 10q arrays have a median CpG density of 9.8 and 9.7%, respectively, the highest in the genome for this window size, and we identified only 8 other genomic regions with densities >7.0%. The next densest region was a chromosome 1 tRNA cluster at 7.9% CpG with a 7.4 kb LM-tRNA repeating unit. This region also displayed methylation gradients that stretch over a ∼50 kb interval. Unlike the D4Z4 arrays, these gradients appear symmetrical on both ends of the cluster. This tRNA tandem repeat has also been shown to contain a conserved CTCF binding site and the repeat unit has features of a genomic boundary element (Darrow and Chadwick 2014). We also observed methylation gradients at the two ends of the chr1 5S rDNA cluster with of repeating 2.2 kb 5S-rRNA repeat, although in this case, the gradients were asymmetric. Several regions of the 45S rDNA clusters were in this extreme CpG density range and there was suggestive evidence of methylation gradients at their borders. Notably, both the chr1 LM-tRNA and 5S rDNA cluster have been identified as one of the few genomic regions that are hypomethylated in the presence of heterozygous *SMCHD1* mutations (Mason et al. 2017). This suggests that shared features among these repetitive clusters with extreme CpG densities may be utilizing common processes to link CpG methylation to the formation of repressive chromatin domains.

Several limitations are associated with our analysis of targeted sequence and methylation analysis of D4Z4 arrays. Our technique improves on the yield of targeted D4Z4 reads from prior studies using nanopore sequencing, however, the yield of full-length reads covering the entire D4Z4 array needs further improvement. Our analysis of near-target reads and the D4Z4 gRNA targeted reads revealed high Cas9 cutting efficiency, so further design of guide RNAs is not likely to improve yield. The presence of near-target reads from divergent D4Z4 reads throughout the genome did not diminish the yield of on-target reads, since the flow cells were still operating at reduced levels of pore occupancy due to a sub-optimal ratio of adaptered molecules to available pores, even with high DNA loads. Because of the complex and repetitive nature of the target sequences, our technique requires a certain level of expertise in mapping of the target reads to appropriate reference sequences. Furthermore, cost for the MinION flowcells is relatively high. Improved pore chemistry, basecalling, and base modification accuracy are likely to provide additional precision, including sequencing the complementary strand from the same molecule. Optimization for lower cost flowcells such as the Flongle, may also help minimize cost and maximize yield. Our analysis for this study relied on genomic DNA from peripheral blood, it remains to be seen whether DNA samples from other tissues will have methylation patterns consistent with those observed here.

Our Cas9 targeted nanopore sequencing assay provides an important tool for genetic analysis of FSHD related D4Z4 arrays. This technique offers an additional tool for clinical diagnosis that can better quantify array length and methylation status, but can also help to provide additional genomic information for unusual cases including somatic mosaicism, 4q-to-10q translocations, and *cis* D4Z4 duplications which are not uncommon in FSHD clinics (Nguyen et al. 2017; Lemmers et al. 2018; Qiu et al. 2020; Rieken et al. 2021). Furthermore, this tool opens the door to a more detailed methylation analysis, including analysis of the methylation gradient across the array, and methylation patterns within the individual repeats. Here, we show that the chromosome 4q and 10q D4Z4 arrays share a high CpG density and methylation gradients that are seen in only a few sites in the genome including tRNA and rRNA clusters, each with unique regulation of copy number and gene expression (Hummel et al. 2019; Hori et al. 2023). Understanding these unique structures and regulatory mechanisms is especially important in light of emerging therapies including antisFfigense knockdown, targeted re-methylation, and other approaches of *DUX4* transcriptional regulation (Chen et al. 2016; Marsollier et al. 2016; Bouwman et al. 2021; Cohen et al. 2021; Himeda et al. 2021; Rashnonejad et al. 2021).

## METHODS

### Subjects and samples

Subjects with clinical history of FSHD and healthy controls were enrolled after consent to participation under a University of Utah Institution Review Board approved protocol. FSHD1 participants had either confirmed clinical genetic testing or positive family history of FSHD with clinical symptoms consistent with FSHD. FSHD2 participants had pathogenic variants in *SMCHD1* identified by clinical genetic testing. Publicly available Oxford Nanopore files were downloaded from the Oxford Nanopore Open Data set under an Amazon Web Services S3 bucket s3://ont-open-data/cliveome_kit14_2022.05/ for the peripheral blood library prepared from a control subject (C4). Genomic DNA was purified from whole blood using standard techniques provided by Gentra Puregene Blood Kits (Qiagen, cat # 158389).

### Targeted nanopore sequencing

Genomic DNA libraries were generated using the Cas9 Sequencing Kit (Oxford Nanopore Technologies, cat # SQK-CS9109). crRNAs are annealed with tracrRNA (IDT, cat # 1072532) by combining: 1 μL of 100μM of crRNA pooled oligonucleotides (see **Supplemental Table S7** for sequences), 1 μL of 100 μM tracrRNA, 8 μL Nuclease-free Duplex buffer (IDT, cat # 1072570) and heated to 95°C for 5 min. Cas9 ribonucleoprotein complexes (RNPs) were assembled in a 1.5 mL microfuge tube by mixing: 79.2 μL Nuclease-free water, 10 μL Reaction Buffer, 10 μL annealed crRNA/tracrRNAs, 0.8 μL HiFi Cas9 Nuclease v.3 (IDT, cat # 1081060) and incubating at room temperature for 30 min. Genomic DNA was dephosphorylated by incubating 3 μL Reaction Buffer, 6 μg genomic DNA and 3 μL of Phosphatase at 37°C for 10 min in a total volume of 30 μL, followed by 80°C for 2 min. to denature. Cleavage and dATP- tailing of the dephosphorylated DNA was performed by adding 10 μL of the Cas9 RNPs, 1 μL 10 mM dATP, 1 μL of Taq Polymerase and incubating at 37°C for 15 min. in a total volume of 42 μL, followed by 72°C for 5 min. to inactivate the enzyme.

Ligation of sequencing adapters to the genomic DNA was performed by combining 20 μL Ligation Buffer (LNB), 3 μL Nuclease-free water, 10 μL T4 DNA Ligase (LIG) and 5 μL Adapter mix (AMX) in a separate microfuge tube. The adapter mixture was then gently added in 2 successive aliquots to the sample and incubated at room temp. for 10 min. The Cas9 library was purified using AMPure XP (Beckman Coulter, cat # A63881) beads. 80 μL of SPRI dilution buffer was added to the library and bound to 48 μL of the beads at room temp. for 10 min. The beads were washed 2x with 250 μL of either Long Fragment Buffer (p13E-11 and pLAM guide RNAs) or Short Fragment Buffer (D4Z4 guide RNA). After a brief drying the beads were resuspended in either 13 μL (MinION samples) or 7 μL (Flongle samples) of Elution Buffer (EB), incubated at room temp. for 10 min. and transferred to a new microfuge tube.

Cas9 libraries were loaded onto either MinION Flow Cells (Oxford Nanopore Technologies, cat # FLO-MIN106D, version R9.4.1 or FLO-MIN114, version R10.4.1) and run on a MinION Sequencing Device (Oxford Nanopore Technologies, cat # MIN-101B) or loaded onto Flongle Flow Cells (Oxford Nanopore Technologies, cat # FLO-FLG001) and run on a Flongle adapter (Oxford Nanopore Technologies, cat # ADP-FLG001) inserted into the MinION Sequencing Device. The MinION Flow Cells were primed with 1000 μL Flush Buffer (FB) + FLT (Oxford Nanopore Technologies, cat # EXP-FLP002). 12 μL of the Cas9 libraries were combined with 37.5 μL Sequencing Buffer (SQB) + 25.5 μL Loading Beads (LB) and loaded onto the MinION Flow Cell. The Flongle Flow Cells were primed with 117 μL Flush Buffer II (FBII) + 3 μL FLT (Oxford Nanopore Technologies, cat # EXP-FSE001). 5 μL of the Cas9 libraries were combined with 15 μL Sequencing Buffer II (SBII) + 10 μL Loading Buffer II (LBII) and loaded into the Flongle Flow Cell. Data collection was controlled using the MinKNOW software on a Linux desktop computer through a USB 3.0 port as described for the MinION Mk1B IT requirements (Oxford Nanopore Technologies).

### Modified basecalling and alignment

Read-level modified basecalling was performed on fast5 files producing FASTQ and bam output files using the GPU-enabled ONT guppy_basecaller v6.3.8 with the options “-- trim_adapters --trim_strategy dna --do_read_splitting” and the dna_r9.4.1_450bps_modbases_5mc_cg_sup.cfg or dna_r10.4.1_e8.2_400bps_modbases_5mc_cg_sup.cfg config files, unless otherwise noted. FASTQ/A sequences were aligned to the T2T CHM13v2.0 reference genome that included a 200 kb accessory sequence for the 4qB D4Z4 region from the HG002 chromosome 4 sequence. Minimap2 (v2.22) and samtools (v1.12) (Li 2021) was used to generate and process alignments using the options “-ax map-ont” and parsing the output with the samtools -F 2304 flag to remove secondary and supplementary alignments. Alignments were processed with custom scripts that regionally extracted FASTA reads for secondary local alignments to 4q, 10q, and 4qB reference sequence using CrossMatch (http://www.phrap.org/phredphrapconsed.html, implementing the Smith–Waterman alignment) for scoring mismatches and assigning reads to haplotypes.

### CpG methylation analysis

Reference-based modified basecalling was performed using Megalodon v2.5.0 (https://github.com/nanoporetech/megalodon) with guppy v.6.4.2 using the Remora modified base model. The megalodon options included the remora settings as “--remora-modified-bases dna_r9.4.1_e8 sup 0.0.0 5mc CG 1 --mod-map-base-conv m C”, the guppy settings “--guppy-config dna_r9.4.1_450bps_modbases_5mc_cg_sup.cfg”, and the output settings as “--outputs basecalls mappings mod_mappings mods”. Single-read methylation plots were generated with modbamtools (https://www.biorxiv.org/content/10.1101/2022.07.07.499188v1) and averaged methylation levels from the reference anchored base calls were parsed from the Megalodon per_read_modified_base_calls.txt output file. Aggregate modified base counts stored in a modified-base BAM files were converted to bedMethyl files using the modbam2bed tool (https://github.com/epi2me-labs/modbam2bed). All statistical analyses of methylation levels were performed with R version 4.2.1. The linear mixed-effect model analysis used the lme4, lmerTest, and broom.mixed packages with the model equation: fitN <- lmer(met_freq ∼ (RepeatN * Sample) + (1+RepeatN | read), REML = F), where met_freq was the repeat-level methylation value, RepeatN was D4Z4 repeat number (centromeric repeat numbering beginning at 0) and read as the nanopore read name. Kernel smoothing of bedMethyl files was performed with the R package smoothr using the smooth_ksmooth function and plotted with ggplot2. Analysis files for HG002 5mC CpG and other methylation from ONT and HiFi data sets produced by the telomere-telomere consortium were downloaded from the Epigenetic profile link at https://github.com/marbl/CHM13 to https://s3-us-west-2.amazonaws.com/human-pangenomics/index.html?prefix=T2T/CHM13/assemblies/annotation/regulation/.

## DATA ACCESS

The nanopore sequencing data generated in this study have been submitted to the NCBI Sequence Read Archive (SRA) repository (https://www.ncbi.nlm.nih.gov/bioproject/) under the BioProject ID PRJNA940957 (see also **Supplemental Table S8**).

## ACKNOWLEDGEMENTS

We are grateful to Jamshid Arjomand from the FSHD Society and Charles Emerson, Lou Kunkel, and the UMass Chan Wellstone Center for FSHD team for helpful discussions in developing the study. We are grateful for the long history of engagement of FSHD families in Utah and for the work of Mark Leppert and Kevin Flanigan and other Utah FSHD investigators over the past 7 decades. This work was supported by the National Institutes of Health [Wellstone P50 HD060848, NINDS K08 NS097631] and the FSHD Society.

