## Supplemental Figures for "Deciphering D4Z4 CpG methylation gradients in fascioscapulohumeral muscular dystrophy using nanopore sequencing"

**Supplemental Figures**. Deciphering D4Z4 CpG methylation gradients in FSHD using nanopore sequencing. Butterfield et al.

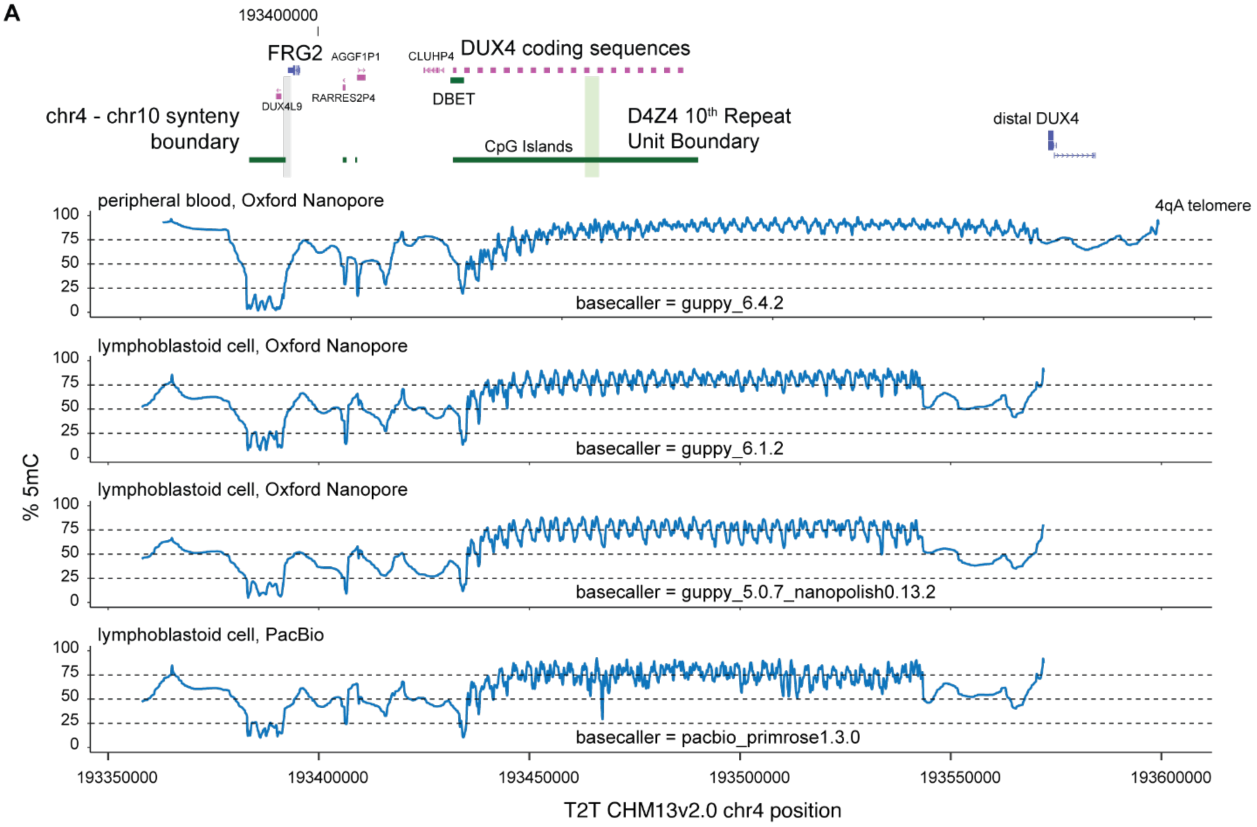


**Figure S1. CpG methylation gradients in the T2T HG002/NA24385 lymphoblastoid sample.** Methylation frequency plots of Oxford Nanopore (ONT) and PacBio HiFi reads. The ONT peripheral blood reads were from the control subject (R10.4.1_e8.2 nanopores) mapped to the 42U haplotype of chr4qA (as in Fig. 2C). The HG002/NA24385 lymphoblastoid cell line data were downloaded from the telomere-to-telomere consortium CHM13 project (https://github.com/marbl/CHM13), using links to the HG002 5mC CpG and other methylation from ONT and HiFi epigenetic profile data. The % 5mC plots were generated from the downloaded T2T bedMethyl files: chm13v1.1_hg002XYv2.7_hg002_CpG_ont_guppy6.1.2.bed, chm13v2.0_hg002_GpC_ont_guppy5.0.7_nanopolish0.13.2.bed, chm13v1.1_hg002XYv2.7_hg002_CpG_pacbio_primrose1.3.0_native.bed.

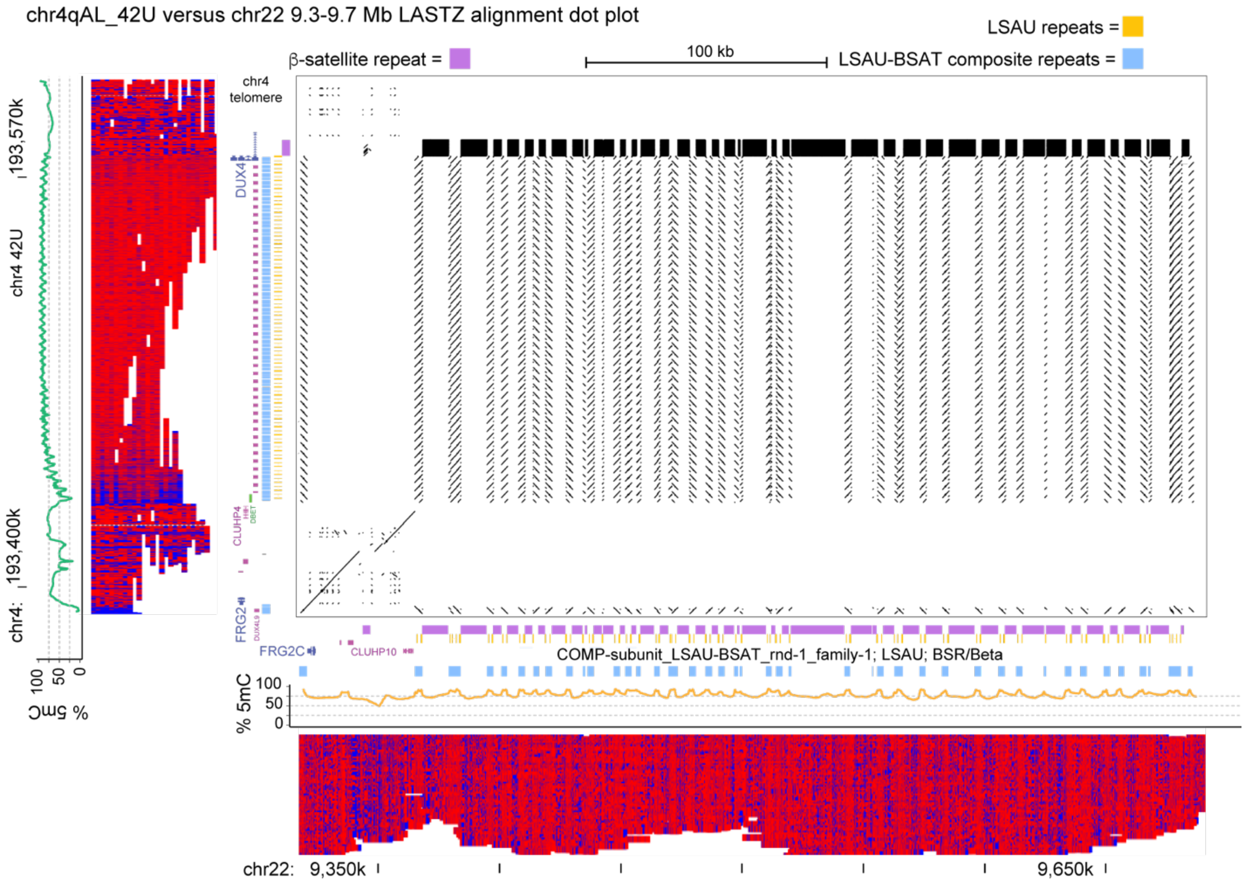


**Figure S2. Paralogous chr22 D4Z4 cluster retains the FRG2 region.**
LASTZ alignment dot plot between the 4qA array on chr4:193,389,177-193,574,945 and the divergent D4Z4 cluster on chr22:9,317,908-9,717,807. The chr4qA sequence has been composited to a length of 41 full-length D4Z4 repeats with a distal 4qAL-pLAM sequence. The data for methylation plots were from participant C4, using Promethion R10.4.1_e8.2 nanopores for sequencing. Modified basecalling was performed with guppy (v.6.4.2) and the dna_r10.4.1_e8.2_400bps_modbases_5mc_cg_sup.cfg config file.

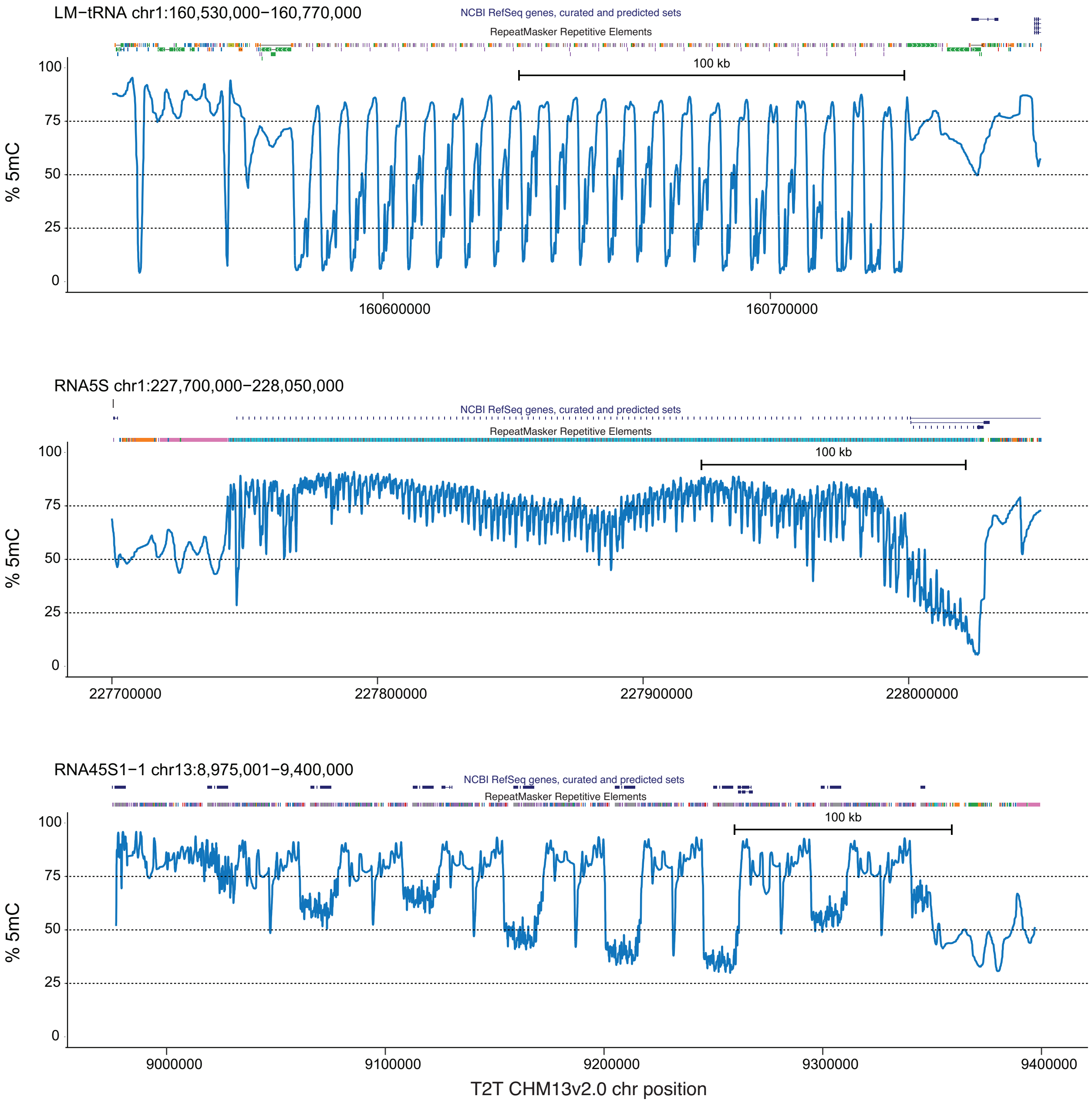


**Figure S3. Methylation gradients in other CpG-dense repeat regions in the T2T HG002/NA24385 lymphoblastoid sample.** Methylation frequency plots of Oxford Nanopore (ONT) reads from the tRNA cluster on chromosome 1, the 5S rRNA cluster on chromosome 1, and one end of the RNA45S1 rRNA cluster on chromosome 13. The % 5mC plots were generated from the T2T bedMethyl files: chm13v1.1_hg002XYv2.7_hg002_CpG_ont_guppy6.1.2.bed.
